# Single Molecule Sequencing of Cell-free DNA from Maternal Plasma for Noninvasive Trisomy Detection

**DOI:** 10.1101/209924

**Authors:** Minyue Dong, Liwei Deng, Huan Jin, Jinsen Cai, Huan Shang, Shuo Zhang, Yueping Zhang, Jinzhou Qin, Dikai Zhang, Feng Long, Renli Zhang, Runsheng Chen, Michael W. Deem, Jun Yu, Jiankui He

## Abstract

The demand of non-invasive prenatal testing for autosomal aneuploidy using cell-free fetal DNA (cffDNA) in maternal plasma is a highly sought-after diagnostic, with a rapidly growing market. Current approaches developed by next generation sequencing (NGS) need PCR amplifcation during sample preparation, which results in amplification bias in GC-rich areas of the human genome. With these approaches, the minimum fetal fraction in maternal plasma is 4% for the small differences in circulating cfDNA between trisomic and disomic pregnancies to be detectable. In this paper, we performed single molecule sequencing of cell-free DNA from maternal plasma for noninvasive trisomy 13, 18 and 21 detections using the GenoCare platform. We found that single molecule sequencing is sensitive enough to detect these chromosome abnormalities when the fetal DNA fraction is as low as 2%. Compared to the Hiseq2500 platform, no significant GC bias was observed. The improved sensitivity and unbiased GC readout make GenoCare a promising platform for autosomal aneuploidy detections, even in the very early stage of pregnancy.

## Introduction

Trisomy 21 (Down syndrome), trisomy 18 (Edwards syndrome) and trisomy 13 (Patau syndrome) are the most common live-birth autosomal aneuploidies. Their incidence of live-birth is as high as 1/700, 1/6,000 and 1/10,000 respectively[1,2]. These diseases can induce serious morbidity and mortality [2]. Amniocentesis and chorionic villus sampling are currently the standard procedures for the diagnosis of fetal aneuploidy [3]. However, these invasive prenatal tests can lead to miscarriage, with rates of total pregnancy loss ranging from 0.6% to 1.9% [4]. To avoid this risk, there is an increasing demand for non-invasive prenatal testing (NIPT) for accurate and safe diagnosis in the early stages of pregnancy.

The discovery of cell-free fetal DNA (cffDNA) in maternal plasma was first reported in 1997[5]. The cffDNA represents a constant relative proportion in maternal plasma [6], thus, sequencing cfDNA opens up a new window for testing fetal genetic conditions. In the first and second trimesters of pregnancy, the faction of cffDNA is 4-10% in total cell-free DNA (cfDNA); and this ratio rises to 10-20% in the third trimester pregnancies [7]. In 2008, the first studies were published showing NIPT of fetal chromosomal aneuploidy by next generation sequencing (NGS) in maternal plasma with Solexa/Illumina platform[9,10]. Cell-free DNA was fragmented in nature, which can be measured by NGS without additional shearing. NGS has been developed as highly sensitive and specific non-invasive screening tools for common fetal chromosome aneuploidies. Detection of fetal trisomy 21 has been reported, but detection of trisomy 13 and 18 seems to be less accurate with this approach [11,12]. The reduced accuracy was due to GC bias influencing sequencing data. A further drawback of NGS is the need for costly equipment and reagents. For example, a trisomy 21 detection test costs roughly $2000 [13].

Amplification-free single molecule sequencing technology can avoid the GC bias of genome. Quake et al. first reported a single-molecule-sequencing technology by a sequencing-by-synthesis method in 2003[14]. Helicos technology, a PCR-free single-molecule-sequencing platform, has been applied for high sensitivity and specificity noninvasive fetal aneuploidy detection without obvious GC bias [15,16]. In our previous study, we demonstrated a direct targeted sequencing of cancer related gene mutations by single molecule sequencing method [17]. Recently, we reported single molecule sequencing of the M13 virus genome and Escherichia coli genome by GenoCare platform developed by Direct Genomics. Here, we presented our study of accurately detecting of trisomy 21, 18 and 13 by single molecular DNA sequencing platform GenoCare.

## Results and discussions

### 1. Sequencing results of trisomy chorinic villus samples

We first demonstrated detecting trisomy positive DNA samples respectively from chorionic villus samples of fetus with trisomy 13, trisomy 18 and trisomy 21 on GenoCare platform in Figure 1. The human genome chromosomes were uniformly segmented into bins containing 150k bases, and then the number of reads mapped back to each bin were counted. The normalized bin depth was defined as the number of reads mapped to a bin divided by the average bin reads of the autosomes. The data is fitted by LOWESS regression as shown in red dots in Figure 1. From Figure 1, we observed that the normalized bin depths of the trisomy chromosomes are significantly higher than those of other normal chromosomes. The Z score is also known as standard score, which is the number of standard deviations between a certain observed value and the mean value of the measured data. Z score is widely used in NIPT. The Z score calculated for chromosome 13 corresponding to Figure 1a is 16.68, that for chromosome 18 corresponding to Figure 1b is 39.98, and that for chromosome 21 corresponding to Figure 1c is 72.39. These results show that GenoCare detects chromosome abnormality in trisomy positive chorionic villus samples with high significance.

**Figure 1.**
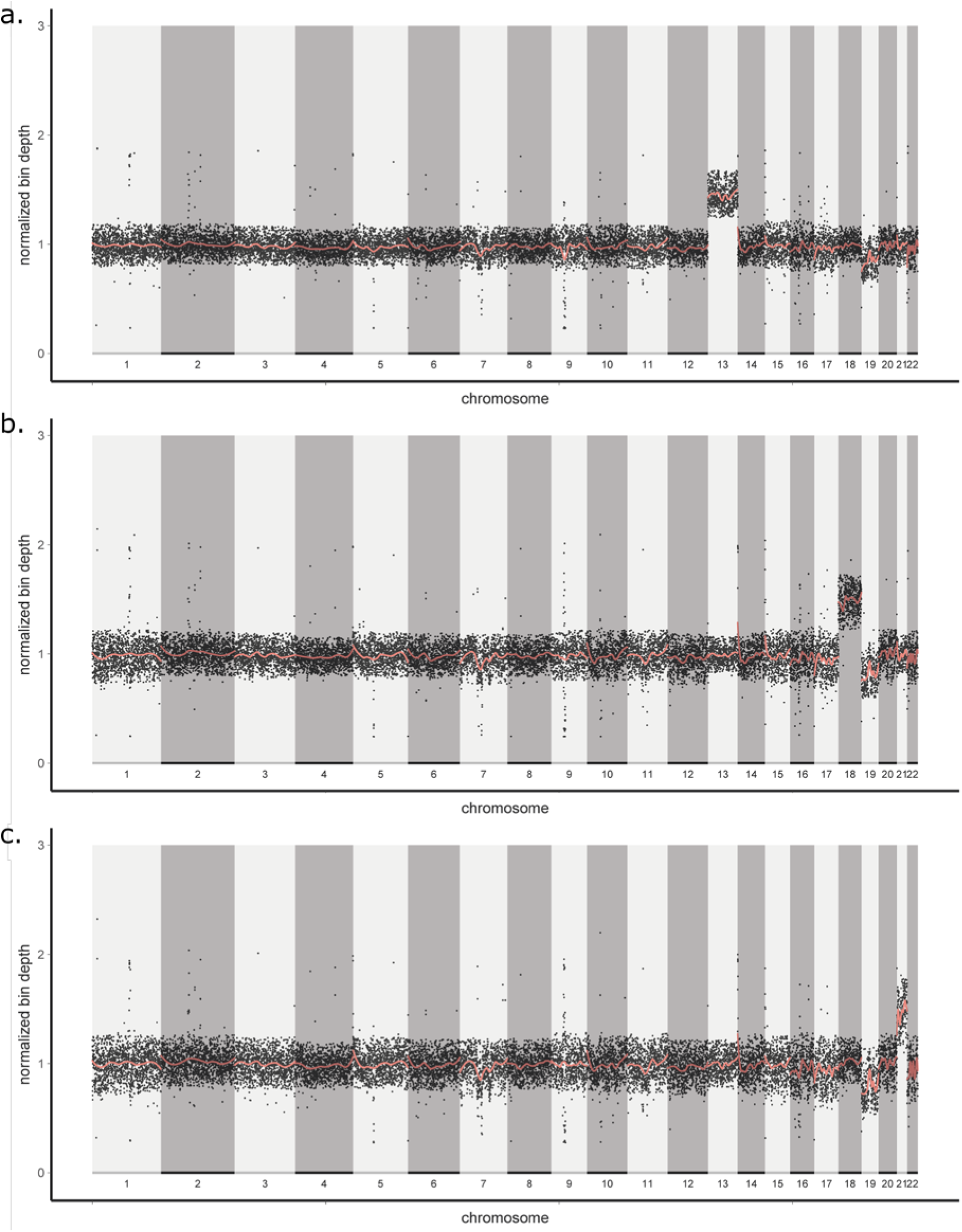
Normalized bin depth at various positions on different chromosomes. The bin size is chosen as 150kbp. **(a),** DNA samples from chorionic villus samples with Trisomy 13. **(b),** DNA samples from chorionic villus samples with Trisomy 18. **(c),** DNA samples from chorionic villus samples with Trisomy 21.

#### Determining the sensitivity by mixing trisomy 21 samples with normal samples

In order to test the sensitivity of the GenoCare platform for NIPT detection, we sequenced samples which were mixtures of genomic DNA extracted from normal women peripheral blood and from chorionic villus samples with trisomy 21. Such mixtures were used to mimic the DNA samples from healthy mother with trisomy 21 fetus. Here we prepared samples containing 2%, 3% and 5% trisomy 21 positive DNA, and the rest part of the samples was normal DNA from healthy people. Figure 2 and 3 present the Z score values calculated for each chromosome from different mixture samples. The Z scores of the chromosome 21 are higher than 3, while the Z scores of other chromosomes are lower than 3. Figure 2,3 shows that the GenoCare platform has successfully detected the trisomy 21 when the DNA mixture contains as low as 2% of DNA from trisomy 21 chorionic villus samples.

**Figure 2.**
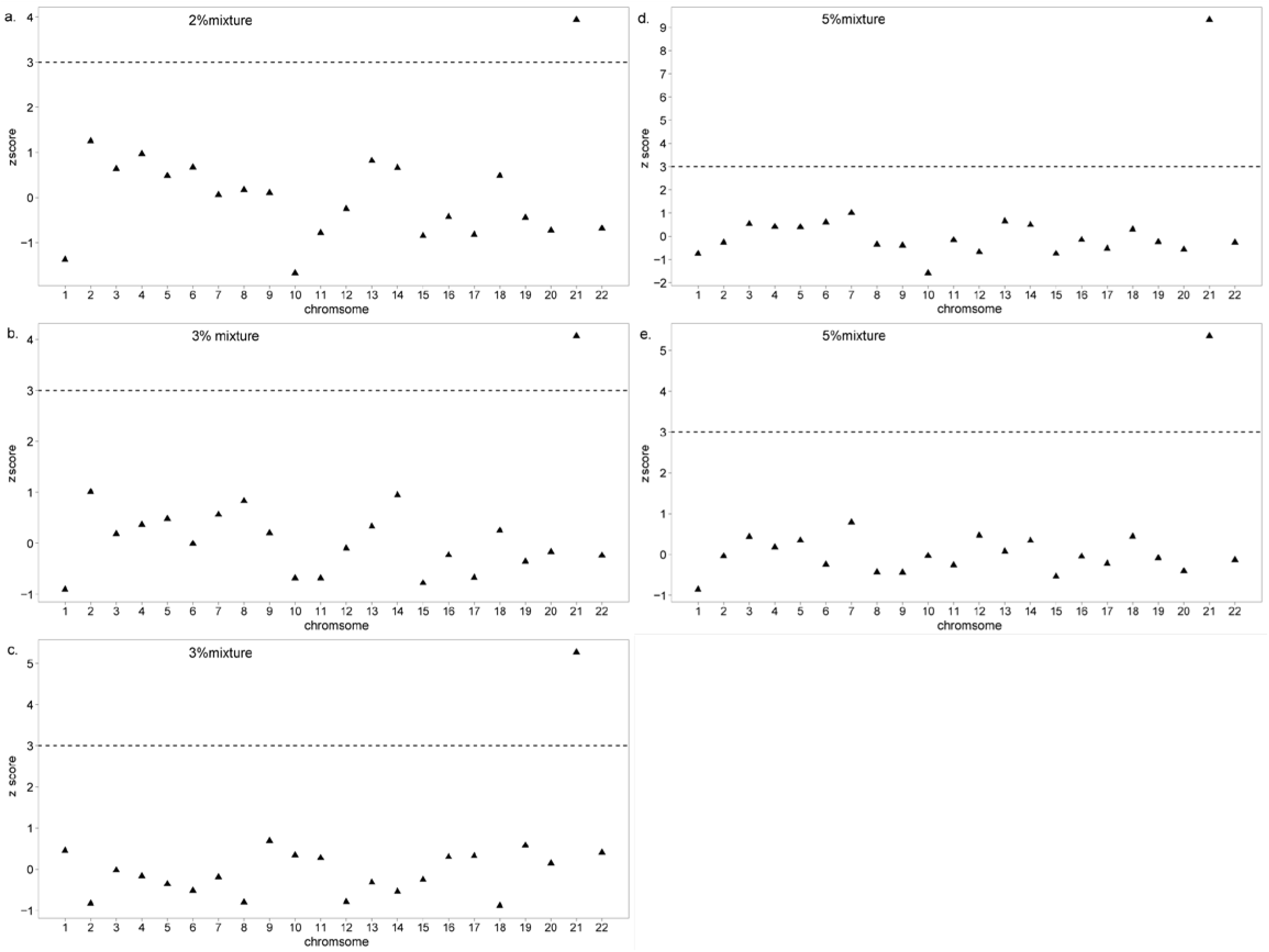
Z scores for each chromosome from samples containing different fractions of trisomy 21 positive DNA. **(a),** Sample with 2% trisomy 21 positive DNA. **(b) and (c),** Sample with 3% trisomy 21 positive DNA. **(d) and (e),** Sample with 5% trisomy 21 positive DNA.

**Figure 3.**
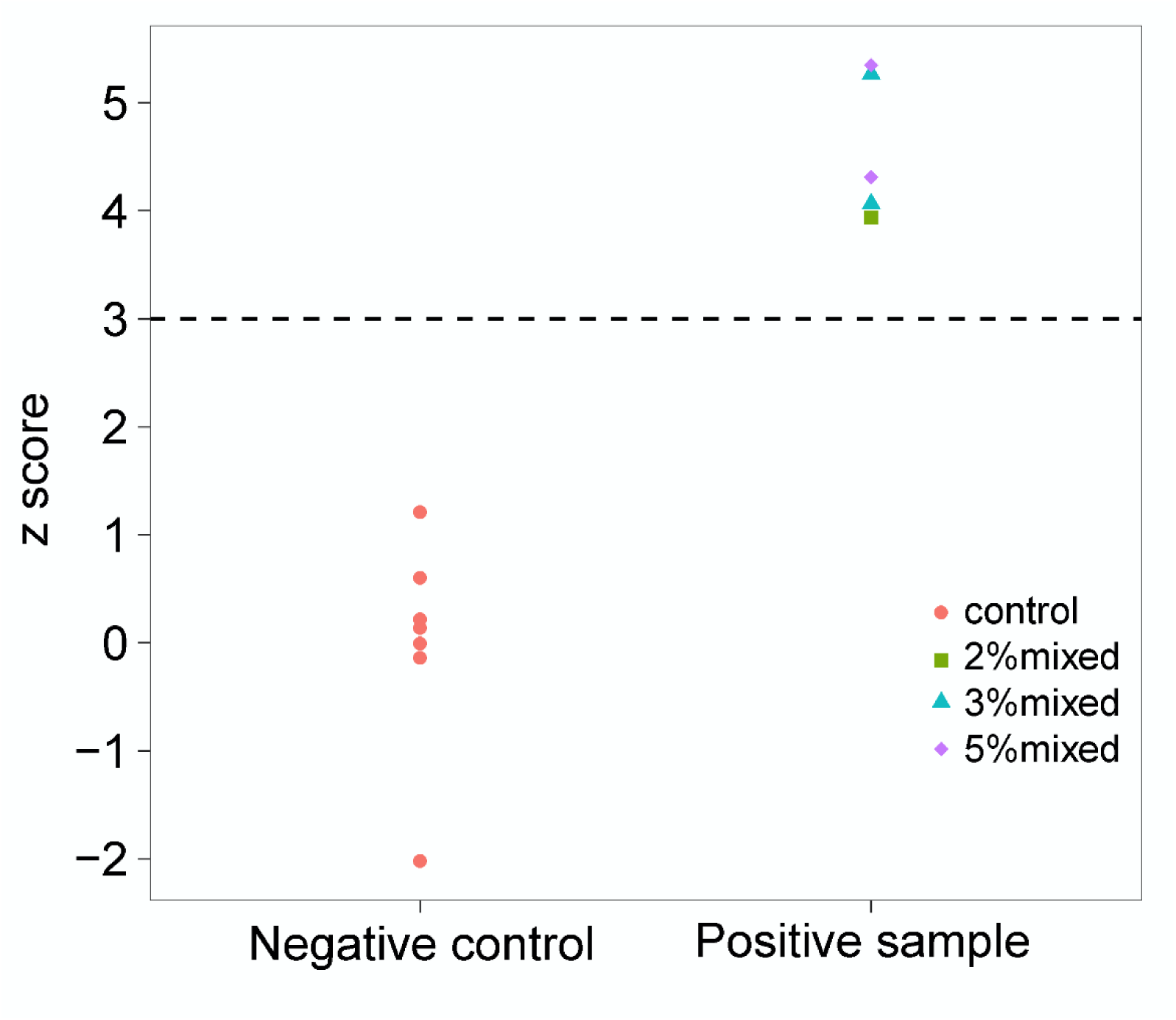
Z scores of the chromosome 21 for samples containing trisomy 21 (positive samples) and for control samples (negative).

#### Sequencing the cell-free DNA sample from maternal plasma

We obtained 7 plasma samples from Women’s Hospital, School of Medicine, Zhejiang University. These samples were collected from pregnant women with 12 weeks gestation, and were confirmed with trisomy 21 by karyotype analysis. We sequenced these 7 samples with GenoCare single molecule sequencing platform. Figure 4 compares the Z scores for chromosome 21 measured for the trisomy 21 positive cfDNA samples and those for negative control samples. The Z scores for chromosomes 21 of the trisomy 21 positive cfDNA samples are prominently higher than 3, and in contrast the Z scores for chromosomes 21 of the negative controls are lower than 3. These results show that we have successfully detected trisomy 21 in seven cfDNA samples.

**Figure 4.**
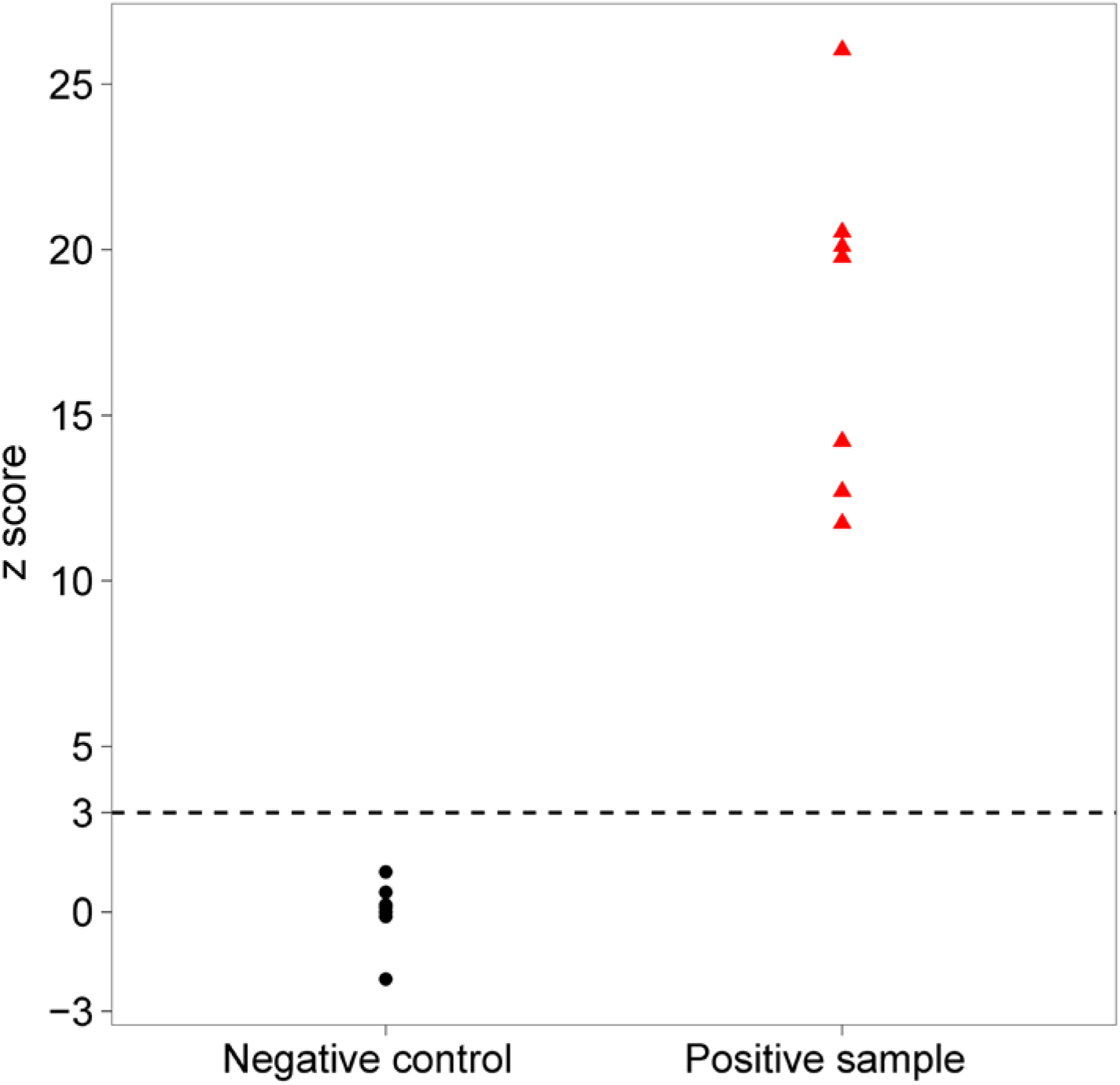
Z scores for chromosome 21 of trisomy 21 positive cfDNA samples (positive samples) and for control samples (negative control).

#### GC bias

A direct comparison of GC bias between the single-molecule-sequencer GenoCare and the NGS platform Illumina Hiseq 2500 was made by sequencing the same DNA samples collected from healthy people on both platforms (Figure 5). The GC bias was calculated as ΔR_GC_^2^, which was defined as the effect of GC bias on the relative reads number [18]. The larger value of ΔR_GC_^2^ represented stronger GC bias. The data recorded by GenoCare showed much less GC bias than Hiseq2500, which can significantly improve the sensitivity for chromosome aneuploidies detection.

**Figure 5.**
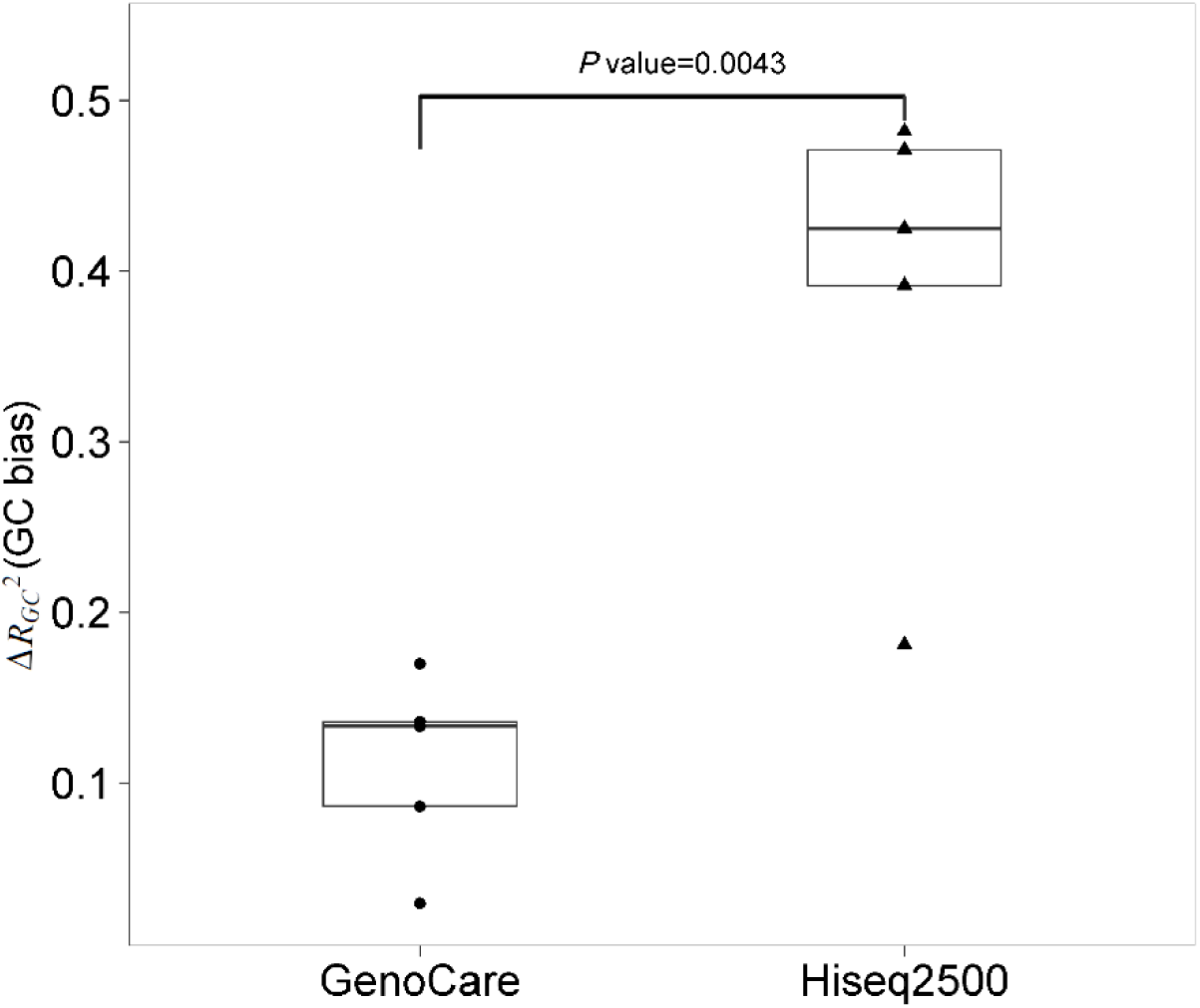
GenoCare platform shows lower GC bias comparing with Illumina Hiseq 2500 platform.

#### Sequencing statistics

The raw reads number, mapped reads number and read length for each positive sample and a representative negative sample were listed in Table 1. The average mapped reads is 5.82 million and the average read length is 34.8bp; The length distribution of the mapped reads for the negative sample presented in Table 1 is in Figure 6. Unlike Illumina or Ion sequencer, GenoCare single molecule sequencing does not suffer from the dephasing issue. The read length is distributed from 28 to 50 bp when the sequencing chemistry was performed in 120 cycles. Longer reads length can be achieved by increasing the sequencing cycles.

**Table 1.** Sequencing statistics on GenoCare platform.

| Samples | Average length of mapped reads (bp) | Genome Coverage | Total reads (Million) | Mapped reads (Million) |
| --- | --- | --- | --- | --- |
| Negative sample | 37.7 | 0.025 | 9.63 | 4.79 |
| Trisomy 21 - 5% mixture | 35.5 | 0.017 | 9.31 | 4.09 |
| Trisomy 21 - 5% mixture | 34.2 | 0.017 | 7.65 | 3.32 |
| Trisomy 21 - 3% mixture | 34.9 | 0.016 | 14.77 | 5.43 |
| Trisomy 21 - 3% mixture | 35.4 | 0.027 | 10.55 | 5.20 |
| Trisomy 21 - 2% mixture | 35.4 | 0.034 | 21.15 | 8.79 |
| Trisomy 21 - 100% positive | 35.2 | 0.023 | 19.18 | 6.98 |
| Trisomy 13 - 100% positive | 34.0 | 0.023 | 20.74 | 7.20 |
| Trisomy 18 - 100% positive | 35.6 | 0.032 | 10.39 | 5.49 |
| Trisomy 18 - cfDNA | 31.6 | 0.010 | 34.78 | 8.06 |
| Trisomy 21 - cfDNA | 33.3 | 0.016 | 13.35 | 4.72 |
| Average | 34.8 | 0.022 | 15.59 | 5.82 |

**Figure 6.**
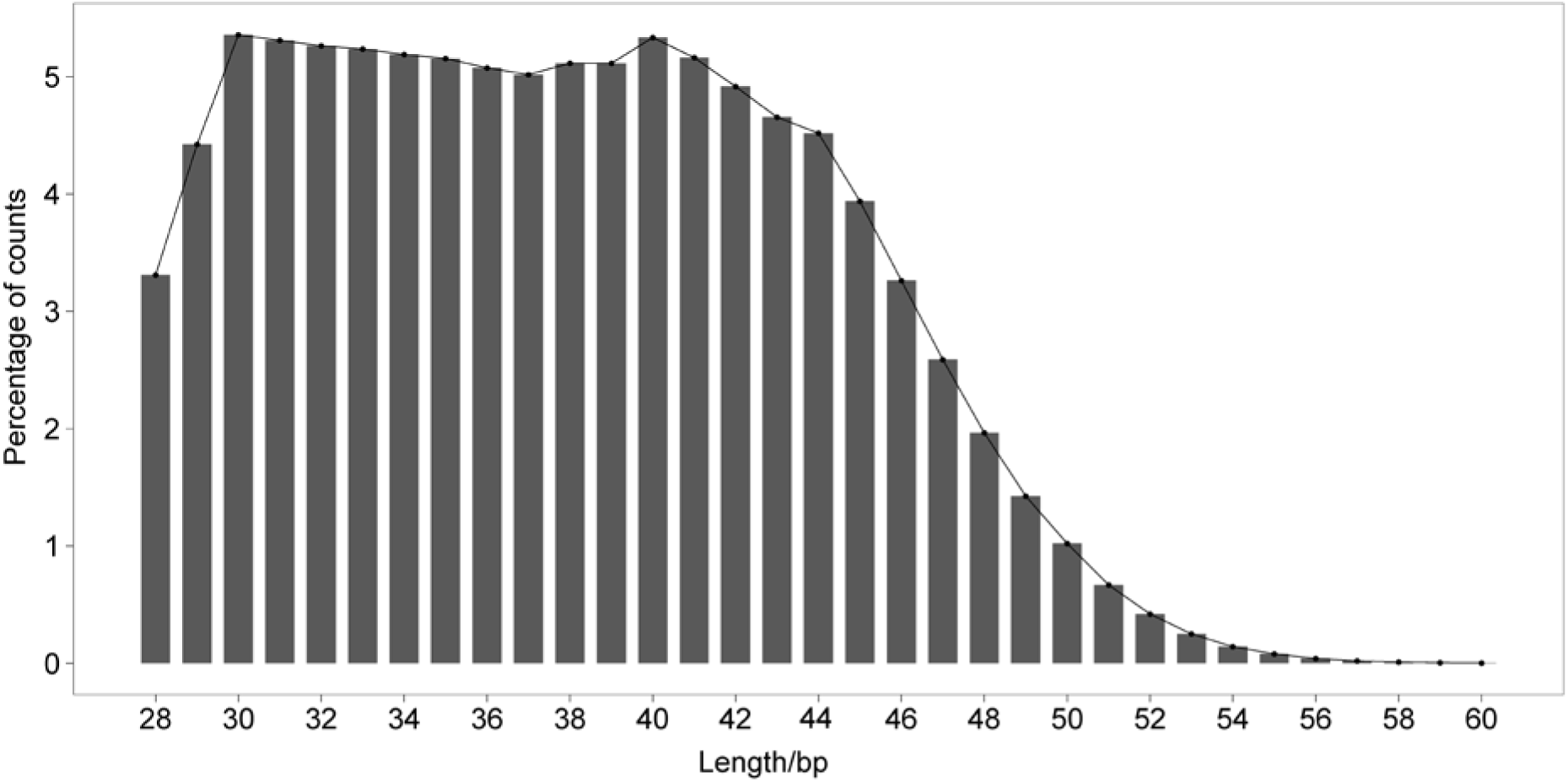
Length distribution of mapped reads for a representative negative control sample.

For the application of NIPT, the ideal sequencing platform should offer enough quantity of valid reads which can be uniquely mapped back to the human genome, while keeping the sequencing time and expenses as low as possible. The read length from the sequencing is a crucial parameter for the subsequent mapping process. Reads with too short lengths are less likely to be uniquely mapped back to the reference genome, while longer read lengths require more base incorporation cycles and thus higher expenditure of time and reagent. Therefore there needs to be a compromise between longer read length and lower cost of time and reagents. Here we studied the relation between unique mapping rate and read length, using GenoCare sequencing data of genomic DNA samples from healthy human. Figure 7 presents the result analyzed using both the DirectAlignment software which we developed in-house and the Bowtie2. The curves show that for the Bowtie2 analysis, the increase of read lengths above 28 bases does not lead to significant improvement in unique mapping rate; for the DirectAlignment analysis, the increase of read lengths between 28 bases and 33 bases results in notable increase of unique mapping rate, however such trend becomes moderate above read length of 34 bases. According to Figure 7, the average of around 35 bases read length we acquired from 120-cycle GenoCare sequencing is good enough for mapping for NIPT purpose, and therefore more sequencing cycles are not crucially necessary.

**Figure 7.**
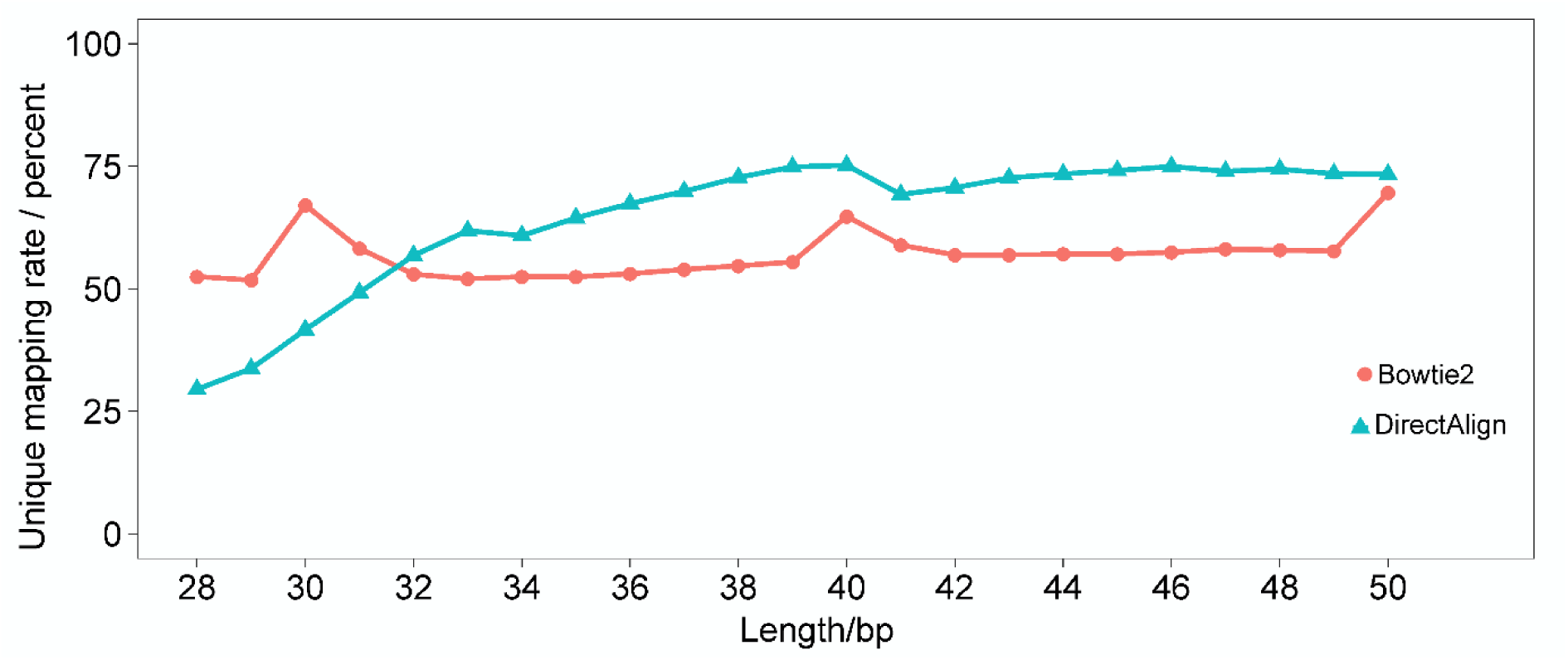
The uniquely mapped reads ratio against read length.

## Conclusion

In this study, we have demonstrated the use of the GenoCare single molecule sequencing platform to detect fetal aneuploidies trisomy 13, trisomy 18 and trisomy 21 with high accuracy. GenoCare can detect the small amounts of trisomy positive DNA fraction as low as 2%. In addtion, comparison between different sequencing platforms revealed that GenoCare platform has less GC-bias compared to other NGS platform. The high sensitivity of GenoCare platform may enable detecting trisomy as early as 8 weeks gestation.

## Materials and Methods

### Ethics and sample collection

This study was approved by the ethic committee of Zhejiang University Affiliated Hospital, and informed consent was received from all patients. Human placental chorionic villi of fetuses with probable chromosome abnormalities and plasma from pregnant women suspected of having fetuses with chromosomal disorder were collected from Women’s Hospital School of Medicine Zhejiang University. All these materials were approved for research purposes in compliance with national guidelines.

### Sample treatment

Genomic DNA (gDNA) of fetus was extracted from placental chorionic villi *via* DNeasy Blood&Tissue Kit (from Qiagen, ref. 69504); Cell free DNA (cfDNA) of pregnant women was extracted from plasma by Magmax cell-free DNA ISO kit (from applied Biosystems by Thermo Fisher Scientific, ref. A29319); gDNA of individual with Down’s syndrome was extracted using QIAamp mini kit (from Qiagen, ref. 51304). In this study, fetus’s gDNA and cfDNA of pregnant women was sequenced, analyzed and compared with the reference sequence to identify if there were chromosomal abnormalities. The gDNA of individual with Down’s syndrome was mixed with gDNA of a healthy female in a ratio of 1:99, 2:98 and 3:97 as a simulative positive sample, to sequence, analyze and compare with the reference sequence.

### Sample preparation

#### 1. DNA fragmentation

The fetus’s gDNA and the mixed gDNA with 1%, 2% and 3% gDNA of individual with Down’s syndrome was randomly fragmented into 200bp dsDNA fragments using WGS-IT fragmentation Enzyme mix (from Enzymatic, ref. SL052517). Agencourt AMPure XP beads (from Beckman, ref. A63881) were then used for fragments purification. The concentration of DNA fragments was assessed by UV absorption using a Nanodrop 2000 device.

#### 2. Poly-A tailing and blocking

Multiple incorporations of 50-100 dATP at the 3’ end of single strand cfDNA from pregnant women and ssDNA fragments from fetus and mixed simulative sample resulted in a poly-A tail. This reaction completed within 20 minutes. In a second step, poly-A tailed 3’ ends were blocked by Cyanine 3 dideoxy ATP (Cy3-ddATP from PERKINELMER, ref. NEL586001EA). The blocking reaction completed within 30 minutes using the enzyme Terminal transferase (from NEB, ref. M0315) such that the incorporation of reversible terminators at the 3’ end of the template strands was prevented.

### Base calling

The base calling method applied in this study is exactly the same as that we reported in previous work [19].

### Mapping method

For the mapping of single molecule sequencing reads, a new software named DirectAlignment has been developed in-house. DirectAlignment performs the mapping using a 3-step strategy.

First, global alignment is carried out to locate the candidate mapping sites on a reference genome for each reads. We adopted both the k-mers hashtable and Burrows-Wheeler-Transform (BWT) algorithms to build a reference in Pre-Alignment. To accelerate the mapping process, all reads were transformed into a binary sequence. With all candidate sites in hand, a fast bit-vector algorithm for approximate string matching based on dynamic programming[19] were applied for filtration. The candidate sites which fail to fulfill the maximum mapping error tolerance were discarded at this stage. After filtration, only a limited number of candidate sites are left. For further precise analysis of substitution, deletion and insertion errors, the Smith-Waterman local alignment algorithm or Needleman alignment algorithm were applied on reads and reference segment. Our aligner used SSE2 to accelerate the edit distance computation. In addition, in order to guarantee a good accuracy of mapping sites, a maximum error tolerance filter and minimum length filter were applied for final mapping results. A read that failed to meet the maximum error tolerance was cut from both two ends base by base until it meet the error tolerance and was accepted as a mapped read or until its length fell below the minimum length requirement and was discarded.

### Calculation of normalized bin depth

The human genome chromosomes were uniformly segmented into bins containing 150k bases, and then the number of reads mapped back to each bin was counted. The bins containing fewer than 10k unknown reference sequences (marked as N) and with greater than 10 bin reads were kept for subsequent processing. The selected bins were corrected for their GC-content using LOWESS fitting method. The number of reads in each bin was divided by the average number of bin reads of the autosomes, and such result was defined as the normalized bin depth.

### Z score

Z score is also known as standard score, which is widely used in NIPT. It is the number of standard deviations between a certain observed value and the mean value of the measured data. Z score is calculated as 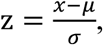, in which *x* is the observed value, *µ* is the mean value of the observation, and *σ* is the standard deviation of the observation.

### GC bias

Here the GC bias is evaluated using ΔR_GC_^2^, which is defined as the effect of GC bias on the relative reads number. ΔR_GC_^2^ was calculated according to the equation 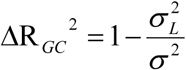[18], in which 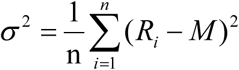 and 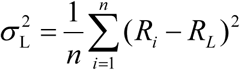. Here *R*_*i*_ refers to the normalized bin depth of each bin, *M* is the global average of the normalized bin depth on euchromosomes, and *R*_*L*_ is the optimal prediction of the normalized bin depth obtained *via* a LOWESS regression fitting of the normalized bin depth against the GC content. Note that here the segmentation of bins on genome is exactly the same as described in previous sections, and each bin contains 150k bases.

## Acknowledgement

We thank Siyu Liu, Yongyi Lu, Yan Li, Lichun Liu, Bo Zhou, Wuxing Liu, Wenzheng Luo, Guanjie Xu, Nannan Zhao, Yuxiang Chen, Bo Yang, Jie Jian, Linsen Li, Weibin Xu, Zengding Wu, Lidong Zeng, Gailing Li, Bin Liu, Weiyue Chen, Zhi Zhao, Fang Chen, Ping Wu, Jia Su, Yan Gao, Luyang Zhao, Zhiliang Zhou, Qin Yan from Direct Genomics for their input and useful discussions. We also thank J. William Efcavitch for his useful advice.

